# Activity-dependent homeostatic synaptic plasticity widens the temperature range of synaptic transmission

**DOI:** 10.64898/2026.08.10.743936

**Authors:** Renato Filogonio, Delaney Cannon, Nikolaus Bueschke, Joseph M. Santin

## Abstract

The framework of homeostatic plasticity posits that neurons regulate cellular properties through feedback homeostasis to maintain activity during changes in the environment. However, when disturbances occur in wild animals they are often caused by environmental variables that induce their own acclimation effects, making it difficult to discern if activity-sensitive feedback plays a role in ecological settings. We addressed this problem using a natural activity perturbation, where frogs hibernate in cold water, leaving brainstem motor circuits that generate breathing inactive for long periods. We show here that motor inactivity, amid complex environmental variables in the hibernation environment, represents a key signal for increasing AMPA-glutamate receptors (AMPARs) on motoneurons. Homeostatic upregulation of AMPARs do not regulate neural activity *per se* but instead correspond with enhanced evoked transmission selectively at cool temperatures. The results show how homeostatic synaptic plasticity may allow animals to restart motor behavior after chronic inactivity encountered in the natural environment. More broadly, these results introduce the concept of homeostatic plasticity as a mechanism to shape thermal tolerance ranges of neural performance in ecological settings.

## Introduction

Animal behavior requires the nervous system to produce electrical activity in the correct amount and timing. This is not a trivial feat, as many environments that animals inhabit are inherently destabilizing. For example, changes in temperature, salinity, and pH alter neural activity and lead to catastrophe at extremes ^1–4^. To explain how neural systems maintain stability in these cases, the concept of homeostatic plasticity provides one framework ^5^. In this view, when activity strays from a set-point during a perturbation, activity sensors transduce that a change has occurred and initiate a suite of compensatory mechanisms involving neuronal modifications that recover activity towards the set-point ^6–9^. While several processes may contribute to environmental stability of neural circuits ^4,10^, negative feedback homeostasis provides an attractive explanation as to how the nervous system maintains healthy function in ever-changing ecological settings.

Although promising, this hypothesis is complicated by the fact that environmental variables elicit their own effects on organisms, making it difficult to link activity sensing in neural circuits to the regulation of circuit activity ^11,12^. For example, temperature acclimation involves heat shock proteins, altered metabolism, endocrine regulation, and systemic physiological adjustments, all acting in tandem to improve physiological performance at the acclimated temperature ^13,14^. Equally obscure are the consequences of compensatory adjustments on neuronal function in changing environments. While homeostatic alterations were originally proposed to regulate firing rate and synaptic transmission ^15,16^, the reach of compensatory responses may be far greater. In particular, the modified ion channel and receptor profile brought about by homeostatic compensation induces robustness to subsequent stressors that were previously not tolerated before compensation occurred ^17–19^. Therefore, the outcomes of homeostatic compensation when assessed over a range of environmental conditions are difficult to predict. Together, how activity-dependent homeostasis manifests in ecologically-meaningful contexts remains an open question.

Here, we address this problem using an animal that endures a large activity perturbation to motor systems as part of its life history. Frogs from northern latitudes hibernate in ice-cover ponds ^20^. While they maintain locomotor capacity and sensory processing, brainstem circuits that produce breathing are completely silent because skin gas exchange maintains aerobic metabolism ^21–23^. Since prolonged inactivity of motor behaviors is a rare trait in vertebrates, especially within the rhythm generating circuits for breathing, this presents an opportunity to address mechanisms of activity-dependent homeostatic plasticity in an ecologically-relevant setting ^24^. Previous work demonstrated a well-studied homeostatic mechanism occurs within motoneurons during hibernation ^25^, whereby AMPA-glutamate receptor function is upregulated, termed synaptic scaling, as seen in various models of activity or sensory deprivation ^7,26–29^. While synaptic scaling is traditionally viewed as an activity-dependent form of plasticity, the degree to which it arises due to inactivity or temperature acclimation in the hibernation environment remains an open question. Indeed, along with upregulated synaptic strength, motor output of breathing improves performance at cooler temperatures following hibernation ^30^, suggesting mechanisms aligned with temperature acclimation may also play a role. Therefore, we tested if motor inactivity or the cold-temperature environment drives synaptic plasticity and then assessed its contribution to synaptic performance over a range of temperatures. We show here that synaptic plasticity in the hibernation environment requires inactivity and is not driven by cold temperature acclimation. In addition, these adjustments correspond with enhanced evoked synaptic transmission over a wider temperature range. These results introduce homeostatic synaptic plasticity as a mechanism for shaping the thermal tolerance of neural circuit performance.

## Results

During hibernation in frogs, respiratory motoneurons increase the amplitude of miniature excitatory EPSPs (mEPSPs) carried by AMPA-glutamate receptors. Consistent with previous reports ^25,31^, hypoglossal motoneurons contributing to the respiratory pump and other orofacial behaviors ^32^ upregulate the mEPSC amplitude following 2 weeks in the submerged hibernation environment when assessed at a common temperature of 22°C. The rank-order plot of the mEPSC distributions from control vs. hibernators produces a linear function (Fig. 1C) that can be used to mathematically transform the distribution of hibernator mEPSCs to match the control (Fig. 1D). Introduced by Turrigiano, et al. ^7^, this outcome has been used to suggest that a multiplicative scaling mechanism upregulates mEPSCs ^31,33,34^. The interpretation that a single scaling factor acting at all synapses across the neuron, multiplying their strengths by the slope of the linear function, has been questioned ^35–37^. Our intent here is not to make explicit inferences about multiplicative scaling across all synapses *per se* or to question that some individual synapses may strengthen more than others. Rather, we confirm the similarity between plasticity in mEPSC distribution caused by hibernation and what has been observed during activity perturbation, commonly referred to as “synaptic scaling.”

**Figure 1.**
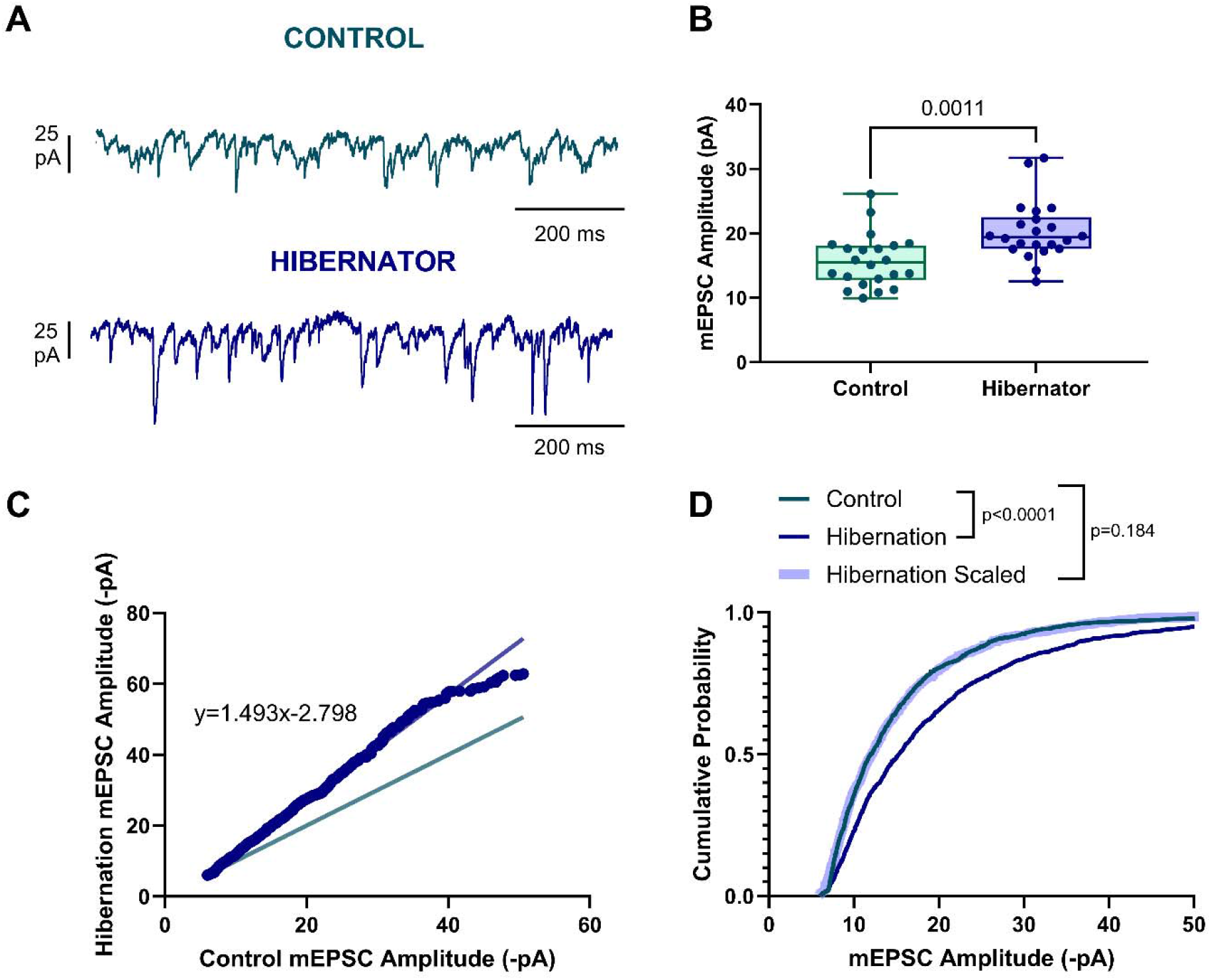
Submerged hibernation increases mEPSC amplitude in inactive hypoglossal motoneurons through a mechanism consistent with synaptic scaling. A. Example of mEPSC recordings from control (top) and submerged hibernator (bottom). B. Box and whisker plots with individual data points from controls and hibernators (control: n=22 neurons from N=8 animals, hibernator n=22 neurons from N=5 animals; t_42_=3.505, p=0.001, unpaired two-tailed t test). C. Rank order of individual mEPSC amplitudes obtained from controls and hibernators. Green line shows unity (ie., if hibernators did not increase relative to controls). Blue regression line is drawn through the plot of control vs. hibernator mEPSCs, where the slope of the line indicates that hibernators increased relative to controls. D. Cumulative probability distribution plots of control, hibernators, and the hibernator mEPSC distribution mathematically downscaled by the linear equation. The hibernator distribution is strongly right-shifted compared to controls, but when downscaling the hibernator mEPSC distribution by the linear equation, hibernators near-superimposed the control distribution (Kolmogorov-Smirnov test; p<0.0001 control vs. hibernator, p=0.1857 control vs. scaled hibernator).

Given that respiratory motor systems are quiescent in the cold hibernation environment ^21^, our goal was to determine if motor inactivity is required to upregulate AMPAR mEPSCs or if a mechanism associated with cold-acclimation drives these responses. To differentiate between these possibilities, we acclimated two groups of animals to ∼4°C. We then removed air access in one group, forcing submergence and therefore respiratory motor inactivity as we have done previously ^25,31^. In the other group, we lowered the water level, allowing activity of the respiratory apparatus throughout the cold exposure. When comparing mean mEPSC amplitudes, submerged animals increased compared to controls; however, mEPSC amplitude in animals with motor activity in the cold (C-MA) matched controls (Fig 2A-B). We observed a slight increase in mEPSC frequency in submerged animals, while C-MA neurons did not differ from submerged animals or controls (Fig. 2C), suggesting an intermediate response that was less apparent in the presence of motor activity. mEPSC alterations were independent from mEPSC kinetics (Fig. 2D-E), passive membrane properties (Fig.2F-G), and intrinsic excitability (Fig. 2 H-I). Collectively, as motor activity in the cold environment prevents or minimizes plasticity of mEPSCs, inactivity is required for synaptic plasticity in the hibernation environment.

**Figure 2.**
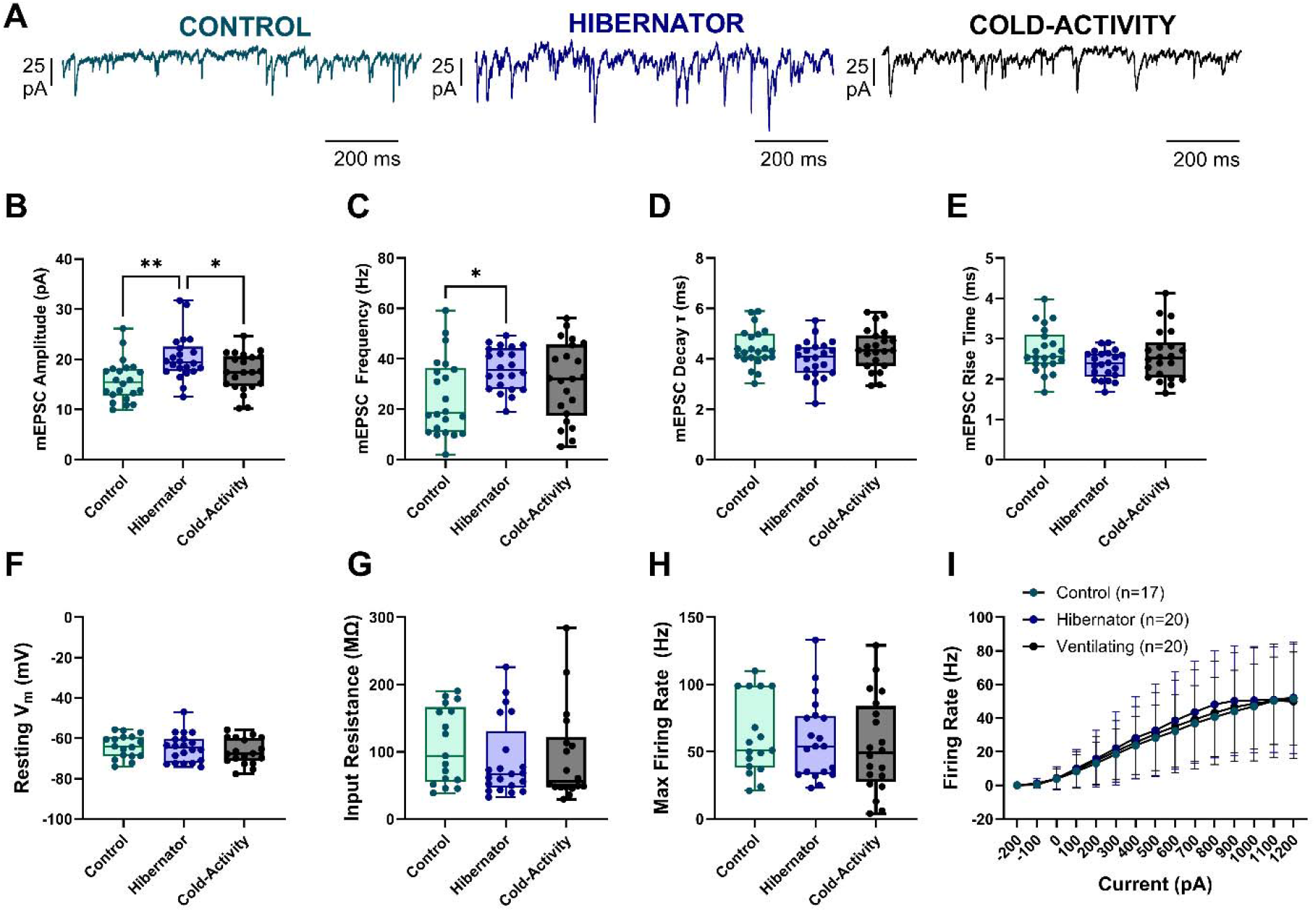
Upregulation of mEPSC amplitude in submerged hibernators requires inactivity and is not driven by cold temperature *per se*. A. Example recordings of mEPSCs from control (left), submerged hibernators (middle), and cold-acclimated animals that maintained motor activity (right). B. Box and whisker plots with individual data points for mEPSC amplitude. Hibernators increased relative to controls, and cold- animals with activity did not increase mEPSC amplitude (one-way ANOVA, F_(2,63)_=6.796, p=0.0021; Holm-Sidak’s multiple comparisons tests: control vs. hibernator=0.0016, control vs. cold-activity=0.1823, hibernator vs. cold-activity=0.493). C. mean data for mEPSC frequency (Brown-Forsythe ANOVA test, F_(2,53.06)_=3.826, p=0.0280, Dunett’s T3 multiple comparison tests: control vs. hibernator=0.0159, control vs. cold-activity=0.4197, hibernator vs. cold-activity=0.5151). D-E. mean data for mESPC decay and rise times (one-way ANOVA for decay, F_(2,63)_=1.944, p=0.1516, for rise time, F_(2,63)_=2.450, p=0.0944). mEPSC data included n=22 neurons from N=8 animal for controls, n=22 neurons from N=5 animal for hibernators, and n=22 neurons from N=7 animals for control-activity. F-G. mean data for resting V_m_ and R_in._ One-way ANOVA for resting V_m_, F_(2,52)_=0.7611, p=0.4723; R_in_, F_(2,53)_=0.5971, p=0.5541. H-I. mean data for firing properties. Maximum firing rate (one-way ANOVA, F_(2,54)_=0.2208, p=0.8026) mean data for frequency-current relationship (no significant group or interaction effects, p=0.8655 and 0.9294, respectively). Passive and active membrane properties data included n=17 neurons from N=8 animals for controls, n=20 neurons from N=7 animals for hibernators, and n=20 neurons from N=6 animal for cold-activity. Data presented as box and whisker plots with individual data points (A-H), and mean±SD (I).

Increases in the amplitude of mEPSCs in submerged hibernators suggest that AMPARs are controlled in an activity-dependent manner, consistent with synaptic scaling ^34^. While synaptic scaling involves the upregulation of postsynaptic receptors, mEPSC amplitude can be influenced by postsynaptic receptor density or neurotransmitter filling within single vesicles ^38^. To address the postsynaptic nature of these responses, we focally applied saturating “puffs” of AMPA to hypoglossal motoneuron cell bodies. To isolate different types of AMPARs, we then blocked Ca^2+^ permeable (CP) forms using 100 uM NASPM, which allowed us to assess 3 currents: the total AMPA-induced current, the NASPM-sensitive current, representing GluR2-lacking CP AMPARs ^39^, and the current remaining in NASPM, representing GluR2-containing Ca^2+^ impermeable (CI) AMPARs (Fig. 3A-B). Consistent with upregulation of postsynaptic AMPARs due to inactivity, the total AMPA-induced current increased in submerged animals, which was also greater than C-MA neurons (Fig. 3C). This pattern appeared to be driven by alternations in both CP and CI AMPARs; however, only CI AMPA receptors were significantly increased in submerged animals compared to controls and C-MA neurons (Fig. 3D-E). Beyond their mean abundance, channels and receptors are known to be co-expressed with quasi-linear relationships that are set by ongoing activity ^9^. Supporting the role of activity-dependent feedback in maintaining receptor co-expression relationships, CP and CI AMPAR subtypes were linearly related in controls (Fig. 3F). The loss of activity in submerged hibernators interfered with this relationship, and no correlation was observed (Fig. 3G), while motor activity in the cold environment maintained the relationship between CP and CI AMPAR (Fig. 3H). Collectively, these results are consistent with central features of activity-dependent feedback that controls the density and coordinated relationships of AMPARs on the background of cold acclimation.

**Figure 3.**
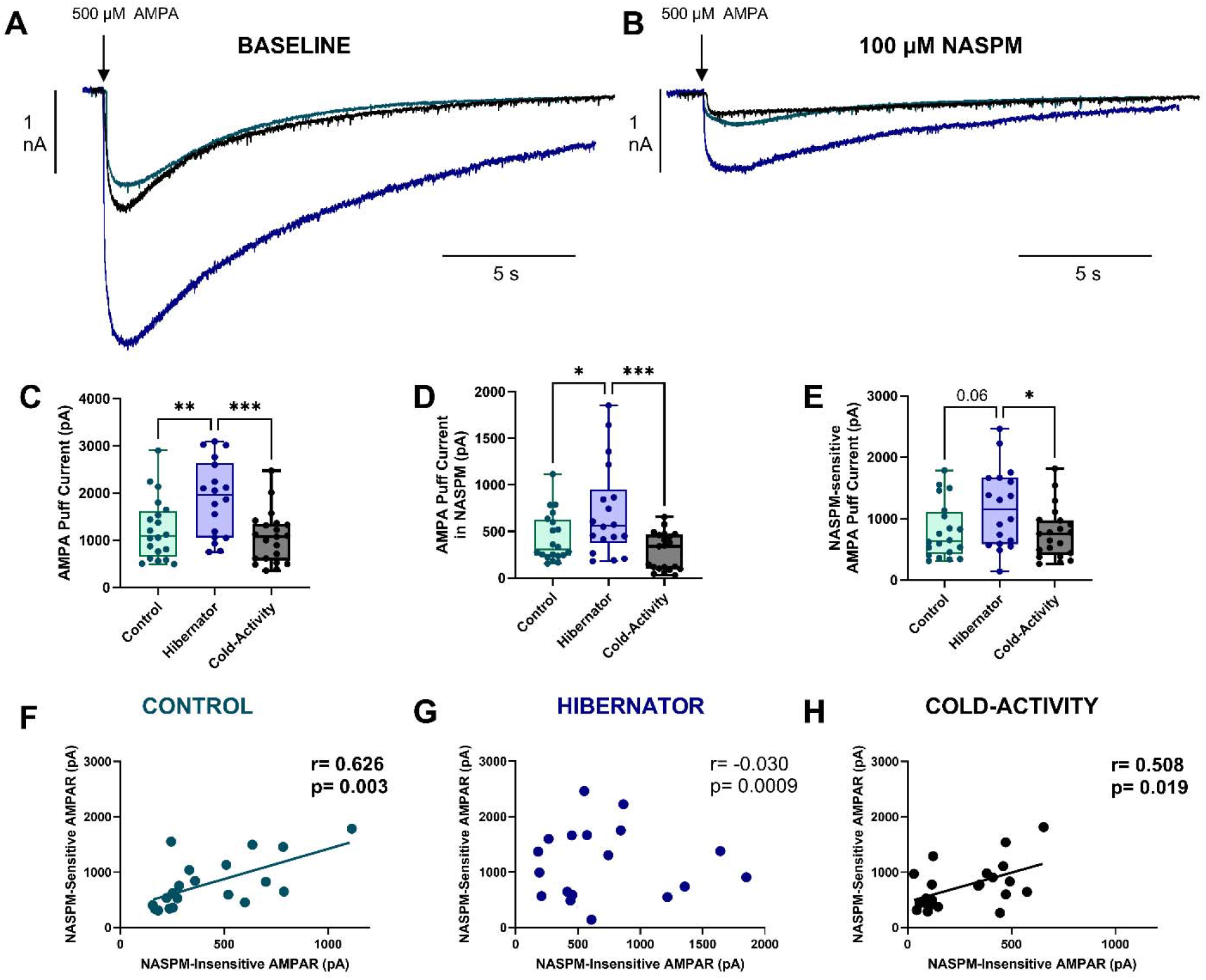
Upregulated postsynaptic AMPA-glutamate receptors involve inactivity but not cold temperature *per se*. A. Example recording of focal application of saturating puff of 500 µM AMPA onto hypoglossal motoneuron cell bodies in control, hibernator, and cold-activity animals. B. These same neurons after bath application of 100 µM NASPM to inhibit Ca^2+^ permeable AMPAR subtypes, with the remaining AMPAR current carried by Ca^2+^ impermeable subtypes. C. Data demonstrating hibernators increase the total AMPAR current compared to controls and cold-activity groups (one-way ANOVA, F_(2,56)_=8.102, p=0.0008, Holm-Sidak’s multiple comparison tests; control vs. hibernator=0.0083, control vs. cold-activity=0.3856, hibernator vs. cold-activity=0.0008). D. Hibernators increase the AMPAR current remaining following NASPM compared to control and cold-activity groups (one-way ANOVA, F_(2,56)_=8.027, p=0.0009; Holm-Sidak’s multiple comparison tests; control vs. hibernator=0.0327, control vs. cold-activity=0.3523, hibernator vs. cold-activity=0.0006). E. The NASPM sensitive current was not significantly greater in hibernators compared to control, but was greater than cold-activity (one-way ANOVA, F_(2,56)_=3.620, p=0.0332; Holm-Sidak’s multiple comparison tests; control vs. hibernator=0.0600, control vs. cold-activity=0.8196, hibernator vs. cold-activity=0.0491). Data in A-C show box and whisker plots with each data point showing data from individual neuron. Data were obtained from n=20 control neurons from N=8 animals from controls, n=18 neurons from N=5 animals, and n=21 neurons from N=7 animals. F-H. Pairwise relationships between the NASPM insensitive AMPAR current and NASMP-sensitive AMPAR currents in controls (F), hibernators (G), and cold-activity (H) groups. Controls and cold-activity showed significant linear relationships between AMPAR subtypes, but submerged hibernators did not. P values and Pearson r are from linear regression analysis.

Activity-dependent synaptic scaling also interfaces with transcriptional machinery that influences the expression of AMPARs ^34^. Using single-cell quantitative PCR, the same design was applied as described above (control, submerged hibernator, C-MA) to measure the mRNA that codes for 4 AMPARs (*Gria1*, *Gria2*, *Gria3*, and *Gria4*) and 2 pore forming kainate receptors (*Grik1* and *Grik2*), each of which may contribute to mEPSCs and/or AMPAR-induced currents (Fig. 4A-F). While expression was variable as is typical in single cells ^9,40^, hypoglossal motoneurons expressed these channel subunits in most neurons. Of these 6 candidate genes, 1 decreased in expression, whereby *Gria3* reduced its expression only in motoneurons from submerged animals. Reduced expression of *Gria3* may seem paradoxical given that mEPSCs and AMPAR currents increased. However, the relationship between mRNA levels and functional expression of the mature AMPAR is often not straightforward in single-cells and are likely to be non-linear due to protein persistence following mRNA transcription and degradation ^41^. These results nevertheless indicate that AMPRAR mRNA is controlled in a manner associated with inactivity and not cold temperature acclimation.

**Figure 4.**
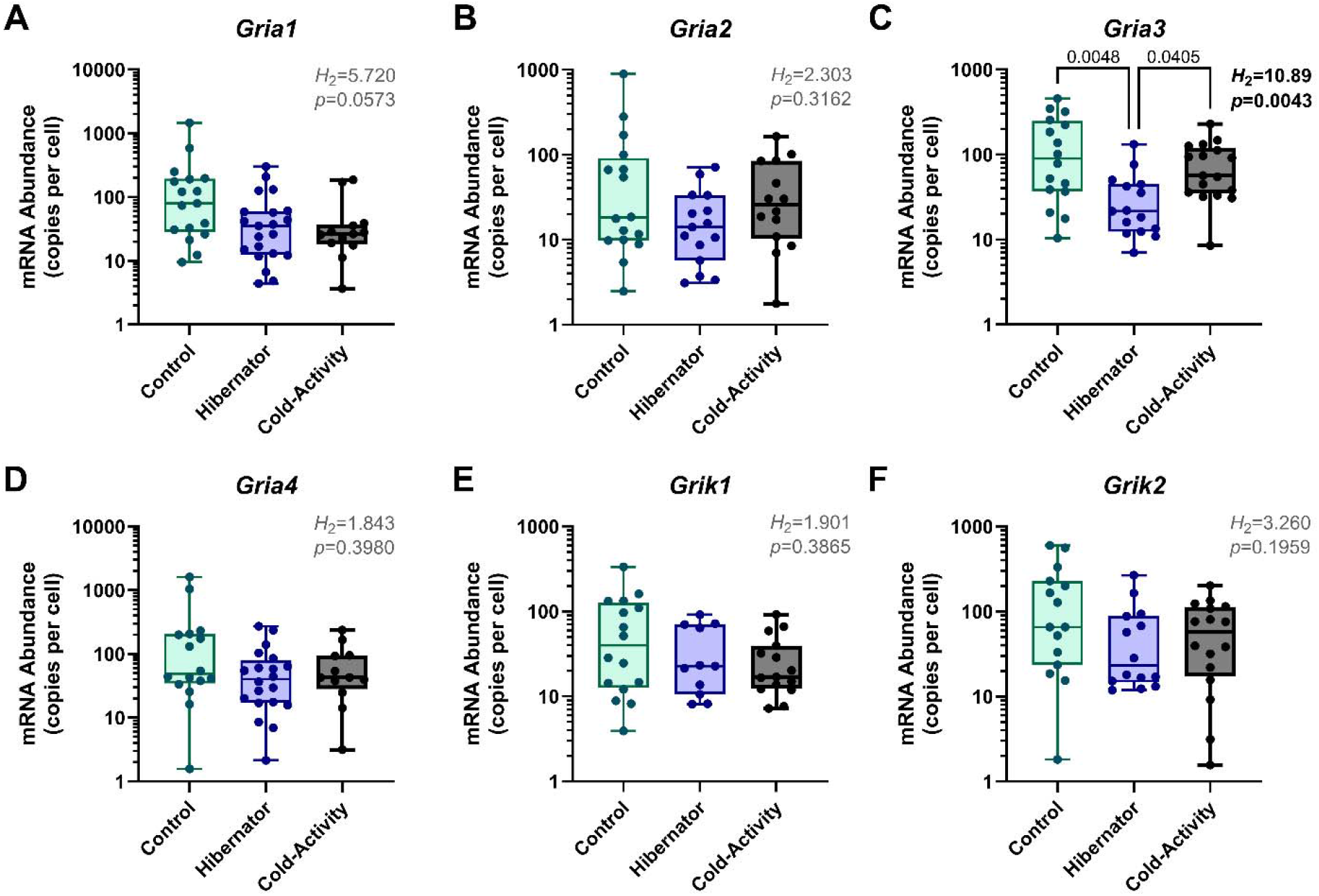
Motor inactivity, but not cold temperature, is associated with single-neuron mRNA changes for glutamate receptors. A-F. Box and whisker plots showing single neuron mRNA counts that code for AMPAR (A-D) and pore-forming kainate receptors (E-F) from controls, submerged hibernators, and cold-activity groups. Only *Gria3* showed a significant difference in hibernators compared to controls and cold-activity groups. All analyzes are Kruskal-Wallis test followed by Dunn’s Multiple Comparisons tests. Data are presented as box and whisker plots with dots representing the mRNA count for one cell.

These results support the hypothesis that synaptic scaling, when embedded in an animal living through an extreme environment, is influenced by activity-dependent feedback at multiple levels. What are the functional implications of upregulated AMPARs? Hypoglossal motoneurons drive contraction of muscles of the buccal floor for lung ventilation and other orofacial behaviors ^42–44^. Since frogs cannot breathe under water, compensatory adjustments that occur during hibernation likely play a role during emergence when motor behaviors restart ^25,30^. Thus, synaptic scaling may lead to greater evoked (i.e., action potential-driven) EPSCs to enhance motor outflow to the respiratory pump, helping to promote breathing after weeks to months under water. In addition, motoneurons of the respiratory network typically suppress their output at cool temperatures, but hibernation induces plasticity that extends the operating range into cooler temperatures. This has also been suggested to play a role to help promote motor activity in the cold upon emergence from the hibernation environment ^30^. To address these possibilities, we measured evoked synaptic transmission (eEPSCs) over a range of temperatures in controls and after activity-dependent synaptic scaling had been induced in submerged hibernators.

For this, we measured evoked excitatory synaptic transmission onto hypoglossal motoneurons through minimum stimulation of solitary tract axons, which play a role in cardiorespiratory functions ^45^. Under these conditions, eEPSCs putatively represent neurotransmitter release due to stimulation of one or a few axons, which in addition to response amplitude also allows for an empirical estimate of neurotransmitter release probability (P_r_) based on the ratio of successful EPSC to total stimulation events ^46,47^. Despite increases in mEPSCs and AMPAR amplitudes (Fig.1-3), eEPSC and P_r_ at putative solitary tract inputs were not different between controls and submerged hibernators (Fig. 5A-C). However, after cooling to 16°C, eEPSC amplitude was larger in hibernators than controls with no change in P_r_ (Fig. 5D-F). At 10°C, the amplitude increased along with P_r_ compared to controls (Fig. 5G-I). Increased synaptic transmission at cool temperatures in hibernators was not associated with altered passive membrane properties compared to controls (Supplementary Figure 1). These results indicate that evoked synaptic transmission is the same at baseline temperature after synaptic scaling has been induced, but greater at reduced temperatures through mechanisms that are independent of P_r_ at 16°C and accompanied by increased P_r_ at 10°C.

**Figure 5.**
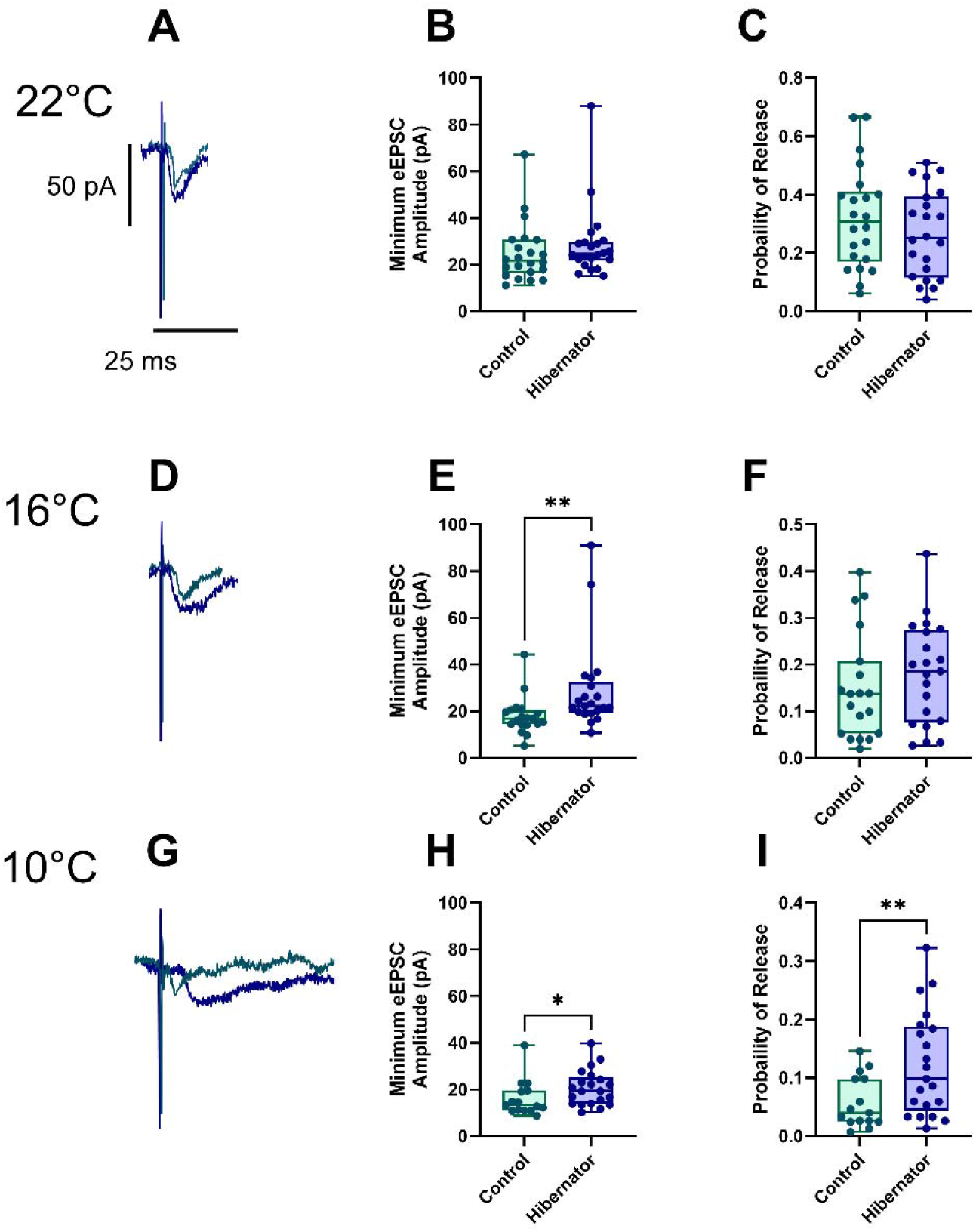
**Submerged hibernators have greater evoked EPSCs at cool temperatures compared to controls**. Evoked EPSCs (eEPSCs) were determined at minimum stimulation. A-C. Controls and hibernators have similar eEPSC amplitudes and release probability (P_r_) at 22°C (amplitude: U=189, p=0.2202, two-tailed Mann Whitney U test, P_r_: t_41.06_=1.082, p=0.2857, unpaired t test with Welch’s correction). D-F. Submerged hibernators have greater eEPSC amplitudes and similar P_r_ at 16°C (Fig. 5D-F; amplitude: U=82.50, p=0.0011, two-tailed Mann Whitney U test, P_r_: U=166.5, p=0.3791, two-tailed Mann Whitney U test). G-I. Hibernators have greater eEPSC amplitudes and P_r_ at 10°C (amplitude: U=91, p=0.0322, two-tailed Mann Whitney U test, P_r_: t_31.03_=2.861, p=0.0075, unpaired t test with Welch’s correction). Control experiments involved N=6 animals and used n=22 neurons at 22°C, 19 neurons at 16°C, and 15 neurons at 10°C. Hibernator experiments involved N=6 animals and used n=22 neurons at 22°C, 21 neurons at 16°C, and 21 neurons at 10°C. Data are presented as box and whisker plots, with dots representing individual data point for one neuron.

We then repeated these experiments at maximal stimulation during temperature ramps from 22°C to 10°C and measured spontaneous EPSC (sEPSC) properties in between evoked events in the same neurons. This allowed us to verify results at minimum stimulation, but also to correlate temperature sensitivity of eEPSCs with spontaneous release amplitude, which relates to synaptic scaling ^31^ and frequency, which is influenced by P_r_ and other presynaptic release properties ^48^. Corroborating results from minimal stimulation, controls and hibernators were the same at warm temperatures, but hibernators had greater eEPSCs compared to controls at temperatures ≤18°C (Fig. 6A-C). sEPSC amplitude increases in hibernators mirrored results from mEPSCs and AMPAR activation, and extended across the entire temperature range from 22°C to 10°C (Fig. 6D). This increase was associated with a group effect but no interaction with temperature in a two-way ANOVA, indicating hibernators had greater amplitudes independent of temperature. This was verified by expressing sEPSC amplitude as a fold change from 22°C, which eliminated the differences across groups at 16°C and 10°C (Fig. 6E). In contrast in amplitude, sEPSC frequency showed a significant interaction between group and temperature in a two-way ANOVA, driven by greater frequency in hibernators at 10°C (Fig. 6F). When expressing sEPSC frequency as a fold change from 22°C, hibernators had greater relative frequences at 16°C and 10°C compared to controls (Fig. 6G), indicating spontaneous release events reduced less (or stayed larger) at cool temperatures than controls. Overall, these results support that hibernators maintained greater transmission at temperatures below baseline, which is associated with enhanced amplitude of spontaneous release events and a greater frequency of spontaneous events at cool temperatures compared to controls.

**Figure 6.**
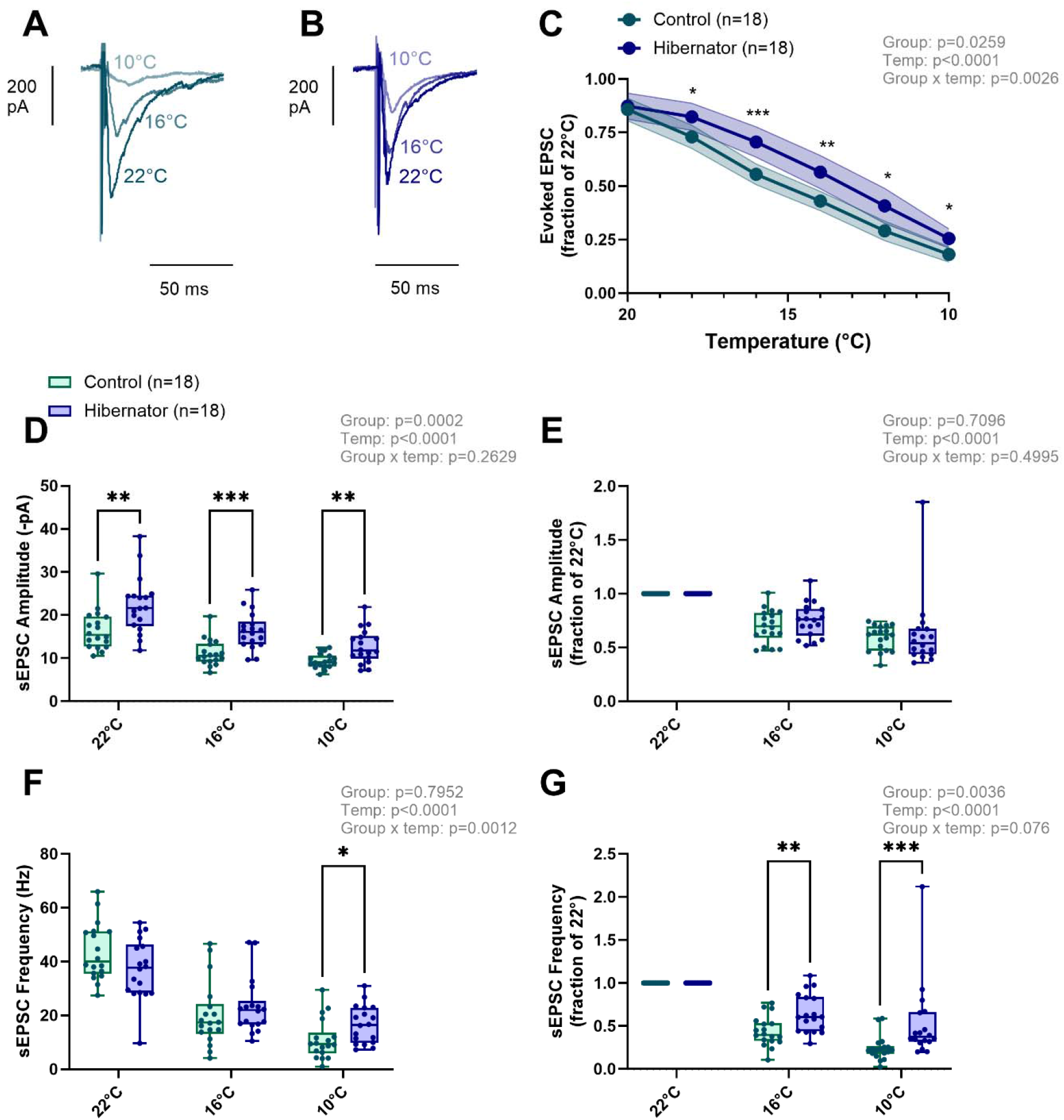
Maximal eEPSC amplitude and spontaneous EPSCs amplitude and frequency are less sensitive to cool temperatures than controls. All data points presented here (eEPSC and sEPSCs) were collected from individual neurons from 22°C to 10°C. A-B. Control experiments involved N=5 animals and n=18 neurons. Hibernator experiments involved N=5 animals and n=18 neurons. Example recordings of maximal eEPSC amplitude from controls (A) and hibernators (B) during cooling from 22°C to 10°C. Hibernators depress less during cooling compared to controls, and therefore, are less sensitive to cool temperature. C. Relationship between maximum eEPSC amplitude normalized to baseline (22°C) and temperature. Hibernators were greater at all temperatures below 20°C. Data are presented as mean±95% confidence intervals. D-E. sEPSC amplitude is greater in hibernators at all temperatures (D). When normalized to 22°C, there is no difference at any temperature, indicating differences at cool temperatures were the result of upregulated baseline amplitude and alterations in temperature sensitivity. F-G sEPSC frequency showed a significant interaction with group and temperatures, indicating altered temperature sensitivity driven by an absolutely greater frequency at 10°C in hibernators. G. shows the data from F normalized to baseline (22°C), demonstrating a greater relative frequency at both 16° and 10°. All analyzes are repeated measures two-way ANOVA followed by Holm-Sidak multiple comparisons test. *** indicates p<0.001, **<0.01, and *<0.05 from Holm-Sidak multiple comparisons test.

To address relationships between eEPSCs at different temperatures and spontaneous release properties, we performed linear regressions between these variables. If synaptic scaling or greater spontaneous release frequency at cold temperatures contribute to enhanced evoked transmission in the cold, we hypothesized that sEPSC properties would correlate with evoked transmission at cool temperatures. In controls, no relationships between eEPSC and sEPSC properties were observed at any temperature (Fig. 7A-F). At 22°C, there was no relationship between evoked transmission and sEPSC amplitude or frequency in hibernators (Fig. 7A-B). However, at 16°C spontaneous release amplitude was proportional to the maintenance of the eEPSC amplitude, whereby neurons with greater spontaneous amplitudes kept evoked transmission closer to baseline (Fig. 7C-D). At 10°C, hibernators maintained the same relationship but also had a significant relationship with spontaneous release frequency (Fig. 7E-F). These results indicate that enhanced evoked synaptic transmission upon cooling in hibernators is associated with greater postsynaptic strength (as indexed by sEPSC amplitude), along with temperature-dependent increases in presynaptic properties (as indexed by sEPSC frequency).

**Figure 7.**
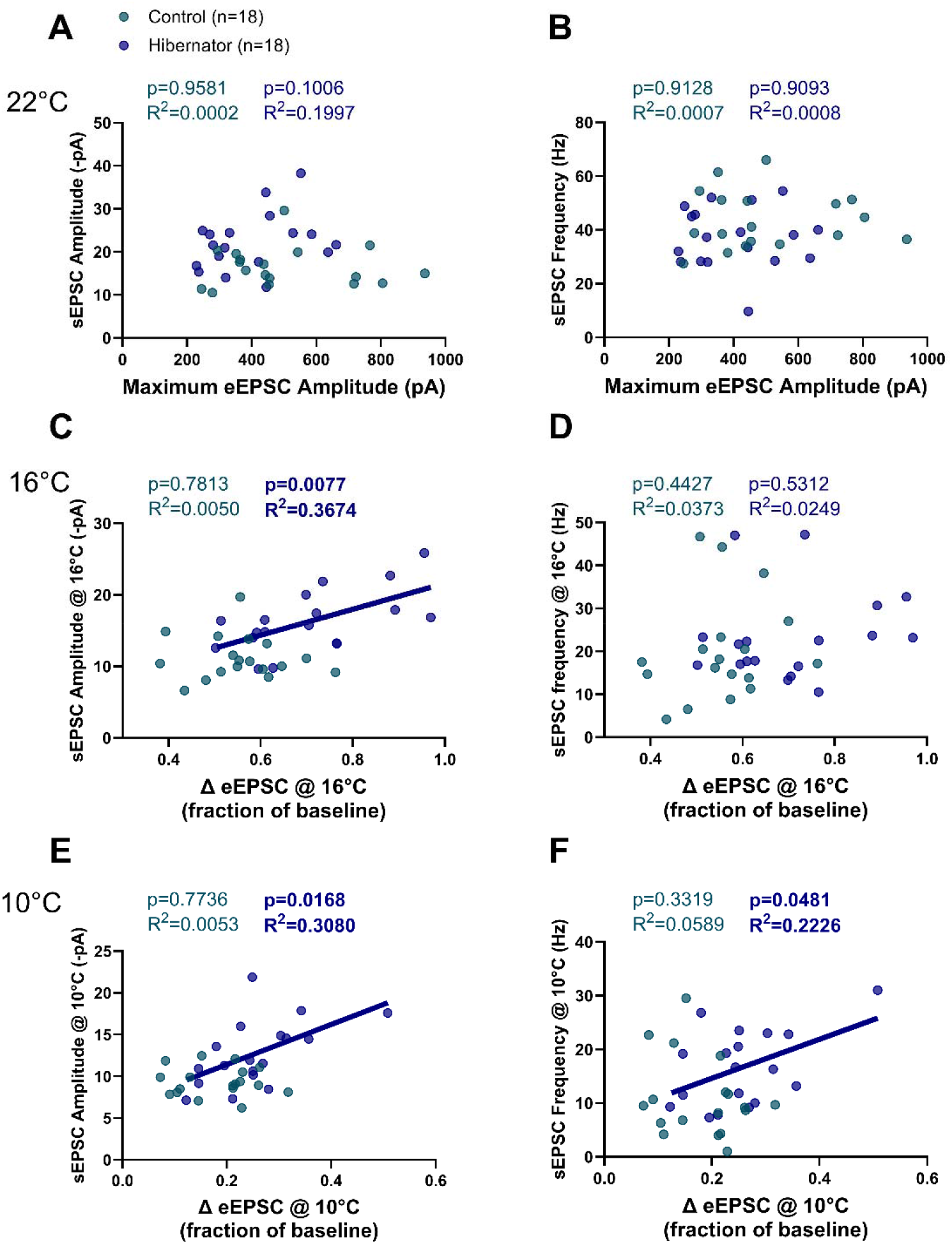
**Improved evoked transmission at cool temperatures correlates with upregulated sEPSC amplitude and frequency**. A-B. Pairwise relationships between baseline eEPSC amplitude and sEPSC amplitude (A) and frequency (B) at 22°C. No relationships exists for controls or hibernators. C-D. Temperature sensitivity of eEPSC (Δ eEPSC, fraction of baseline) compared to sEPSC amplitude (C) and frequency (D) at 16°C. No relationship existed for controls. C shows that hibernator neurons that had smaller reductions in eEPSC amplitude (ie., they were less sensitive to temperature) had greater sEPSC amplitudes. E-F. Temperature sensitivity of eEPSC (Δ eEPSC, fraction of baseline) compared to sEPSC amplitude (C) and frequency (D) at 10°C. No relationship existed for controls. E. shows that hibernator neurons with smaller reductions in eEPSC amplitude had greater sEPSC amplitudes. At 10°C, neurons with smaller reductions in eEPSC amplitudes also had greater sEPSC frequencies.

## Discussion

The most prominent framework for neuronal homeostasis asserts that activity-dependent mechanisms sense and counteract disturbances by altering synaptic and cellular function (i.e., homeostatic plasticity). However, it is not clear how activity-dependent mechanisms interface with acclimation to physical or chemical environments that initially cause activity disturbances in ecologically-realistic contexts. When frogs hibernate in cold water without motor activity of breathing, we find that activity-dependent mechanisms, rather than temperature *per se*, upregulate glutamate receptor function in a compensatory manner. However, when addressing transmission across a range of temperatures, these homeostatic adjustments were associated with improved synaptic performance selectively at cooler temperatures. These results show that homeostatic tuning rules within neural circuits influence the thermal tolerance of brain function, representing a neural mechanism allowing animals to respond to changing environments. Below we describe mechanistic interpretations, as well as areas for future investigation.

### Homeostatic processes upregulate AMPARs during hibernation

One major goal of this study was to assess whether homeostatic plasticity or activity-independent mechanisms aligning with temperature acclimation modifies synaptic strength during hibernation in frogs. Our results point to homeostatic, activity-dependent processes, as cold animals that maintained motor activity of the buccal floor musculature did not exhibit synaptic plasticity. This is supported by several physiological and molecular assessments. Specifically, inactivity but not cold-acclimation upregulates mEPSCs and AMPA-glutamate receptors, which involved increases in Ca^2+^-impermeable and to a lesser extent Ca^2+^-permeable subtypes, as well as a disruption to their linear co-expression set by ongoing activity. Through assessments of single-cell AMPAR mRNA transcript abundance, we suggest these processes may be associated with activity-dependent processes that control mRNA levels, as the message for *GluA3* was reduced only in submerged hibernators. While the connection between single-cell mRNA expression and the specific AMPARs currents remain to be further investigated, reduced AMPAR mRNA may seem at odds with the functional current measurements in these same cells. However, we hypothesize that mRNA may be upregulated early in the hibernation process to drive protein translation but then undergo degradation as to not further upregulate AMPA receptors later on, leaving AMPAR proteins elevated in the membrane ^41^. While our findings draw a line to motor inactivity as the cause of AMPAR plasticity, the specific signaling mechanisms and physiological processes that underly these responses are not yet known. Activity sensing can represent a variety of different cellular processes that include changes in firing rate, receptor activation state, glial signaling, and more ^28,34,49^. In addition, submerged hibernators have lower metabolic rates than animals maintained at cold temperatures with air-access ^50^. Therefore, reduced metabolism may also interface cellular mechanisms induced by inactivity. Regardless of the specific mechanistic implementation, our results align with key aspects of activity-dependent signaling to drive synaptic compensation in a complex physiological environment.

### What is synaptic scaling doing at warm temperatures?

Synaptic scaling is often interpreted as a multiplicative increase in postsynaptic receptors at all synapses ^7^. Therefore, one would expect enhanced evoked transmission at all inputs. Yet, this did not occur when assessed at warm temperatures despite greater mEPSCs and agonist-induced AMPAR currents (Fig.5-6). These results support for idea that all synapses do not strengthen equally during inactivity ^36^. However, it is now appreciated that mEPSCs use different vesicle release mechanisms, active zones, and/or postsynaptic locations than those participating in evoked release ^51^. Thus, synaptic scaling may selectively enhance the size of spontaneous synaptic events without affecting evoked transmission at select inputs, an interpretation that seems plausible given there was no correlation between evoked EPSCs and spontaneous EPSC amplitude and frequency at warm temperature (Figure 7A-B). We have previously suggested that synaptic scaling represents a way to enhance motor output associated with breathing after hibernation ^25^, but this seems unlikely at warm temperatures in hypoglossal motor neurons since evoked transmission was unaffected. Spontaneous release of neurotransmitter vesicles has been proposed to contribute to maintaining structure of the postsynaptic membrane and dendritic protein translation ^51^. Therefore, functional outcomes of boosting spontaneous transmission may be more important than action-potential dependent processes at warm temperatures after hibernation.

### What is synaptic scaling doing at cooler temperatures?

While evoked transmission was unchanged at warm temperatures, the same neurons from hibernators showed greater evoked transmission during acute cooling (Fig. 5-6). Even more striking was the amplitude of spontaneous release events, which underwent the same increase as observed in mEPSC and AMPAR currents, correlated with greater transmission at cool temperatures (Fig. 7). These results suggest that synaptic scaling may present an advantage by enhancing evoked transmission selectively at cooler temperatures. To explain this relationship, we must reconcile how upregulated postsynaptic AMPA receptors presumably at spontaneously release sites do not play a role in evoked transmission at warm temperatures but then contribute at cool temperatures.

Presynaptic terminals have active zones that favor evoked or spontaneous release. However, in some cases, spontaneous and evoked release sites are not fully segregated, but rather, exist as a spectrum with a relatively high and low chance of release through each mode, with postsynaptic receptors isolated to each release site ^52,53^. If synaptic scaling increases AMPARs at sites receiving vesicles with high spontaneous/low evoked release, this would have little influence on evoked transmission as we show for warm temperatures. We hypothesize, therefore, that hibernators shift the balance from spontaneous to evoked release at sites associated with upregulated AMPARs upon cooling. While the specific mechanisms that allow this to occur remain to be determined, alterations in vesicle-associated proteins, calcium sensors, and calcium channel proximity to primed vesicles all increase all the likelihood of action potentials releasing vesicles from active zones that otherwise release spontaneously ^51^. The types and amounts of voltage-gated calcium channels (VGCCs) vary across active zones even within individual nerve terminals, leading to the differential control over vesicle release ^54^. This is especially relevant, as VGCC subtypes vary in temperature sensitivity, where some pass more calcium at cooler temperatures due to strongly reduced inactivation kinetics ^55^. Indeed, hibernators have enhanced release probability and spontaneous release frequency compared to controls at cool temperatures, pointing to alternations in the temperature sensitivity of neurotransmitter release. Therefore, the mechanistic hypothesis that arises from this interesting relationship is that hibernators shift the balance from spontaneous to evoked release at cool temperatures by favoring the expression of VGCCs with temperature-sensitive properties, permitting the release of vesicles contacting postsynaptic membranes with upregulated receptors. Following this line of questioning may provide new insights into how homeostatic mechanisms interface with changing environments.

### Prospectives for the frog

Homeostatic plasticity is often interpreted as a way for neurons to regulate activity around a set point; when activity drops, homeostatic processes alter neurotransmitter receptor density or intrinsic excitability to restore baseline activity ^5^. In hibernating frogs, the motor processes for breathing are silent while the animals are submerged in cold water ^21^. Therefore, homeostatic compensation of motoneurons brought about by inactivity does not restore motor activity in a traditionally homeostatic sense. However, frogs at the bottom of a pond must pass from cold to warm body temperatures upon emergence and restart motor behavior on this background, conditions where synaptic transmission is otherwise strongly depressed. To work around this problem, we identified (in)activity-dependent increases in AMPARs that arise during hibernation correspond with enhanced transmission at cool temperatures. This mechanism may work alongside decreases in GABA_A_ transmission within the rhythmic generating circuits, which also enhance network activity at cold temperatures upon emergence ^30^, and potentially other mechanisms that alter the thermal tolerance of network activity. Therefore, we suggest that synaptic scaling presents the advantage of bolstering transmission in the cold environment that otherwise strongly depresses synaptic performance to help restart motor behaviors when needed.

### General perspectives for neuronal function in a changing world

A fundamental challenge faced by all animals is to maintain some degree of nervous system stability while traversing an ever-changing array of environments that can disrupt nervous system function. Temperature is especially relevant, as all species besides mammals and birds are ectothermic, where body temperature readily changes as function of ambient temperature. Indeed, climate change is not only associated with warming average temperatures, but also a higher incidence of both warm and cold extremes ^56^. Even for mammals considered to be strict homeotherms, certain brain regions undergo physiological fluctuations that span 4°C, influencing network function and behavior ^57^. In this study, we demonstrated that activity-dependent processes are crucial for adaptation to neural activity disturbances induced by the hibernation environment but, surprisingly, corresponded to a widened range of synaptic transmission during acute temperature alterations. These results build upon a growing set of examples demonstrating that homeostatic compensation can have unpredictable effects on tolerance to subsequent acute stressors ^17,18,58^, and introduce a new way that homeostatic synaptic plasticity builds robustness into neural circuits during environmental change. While we addressed this using amphibian hibernation, we assert that contextualizing homeostatic plasticity as an ecologically-relevant concept should draw attention to other environmental challenges and physiological states associated with activity challenges to the nervous system.

## Material and Methods

### Animals

All experiments performed were approved by the Animal Care and Use Committee (ACUC) at the University of Missouri (protocols #39264 and #65623). Adult American Bullfrogs, *Aquarana catesbeiana*, of undetermined sex were purchased from Rana Ranch (Twin Falls, Idaho, USA) and Niles Biological (Sacramento, CA, USA). Controls and submerged hibernators were maintained as previously described ^30,40^. Briefly, animals were kept in plastic tubs with dechlorinated tap water at room temperature bubbled with room air. Control frogs were acclimated to these conditions for at least a week after arrival at the laboratory before experiments and had access to wet and dry areas in the tanks. Pellet food was provided once per week and was eaten ad libitum. Submerged cold frogs were kept in plastic tanks in temperature control incubators for > 1 week before temperature was lowered in stepwise manner to 4°C over 7 days in a walk-in temperature-controlled environmental chamber. Once water temperature reached 4°C, air access was blocked using a plastic screen placed in the tank. Given that synaptic changes occur within 2 weeks in the hibernation environment and are not different at 4 weeks ^31^, experiments commenced after 2 weeks of submergence.

In the experimental group involving cold-acclimation of animals that maintained motor activity of the buccal floor, these animals were cooled at the same time as animals that were submerged. However, upon reaching 4°C, instead of placing a screen at the water level, most of the water was removed from the tank to encourage ventilation of the buccal cavity. Care was taken to ensure the snout of the animal could not sit below the water later. Upon visual inspection during daily animal welfare checks, all animals were observed to perform buccal pumping, which is driven in part by hypoglossal motor neurons used in this study.

## Tissue Preparations for Physiology Experiments

### Slice preparation, whole-cell patch clamp electrophysiology, and data processing

Tissue slices containing hypoglossal motoneurons for patch clamp electrophysiology experiments were generated following previously established protocols ^59,60^. Briefly, frogs were initially anesthetized with isoflurane (1 ml per liter, v/v) within a plastic box of approximately 1 L. After loss of pedal reflexes (Neto et al., 2024), frogs were decapitated with a guillotine and the head was transferred to a cold artificial cerebrospinal fluid solution (aCSF, in mmol^-1^: 104 NaCl, 4 KCl, 1.4 MgCl_2_, 7.5 d-glucose, 40 NaHCO_3_, 2.5 CaCl_2_, 1 NaH_2_PO_4_, gassed with 1.5% CO_2_/98% O_2_ for pH ≈ 7.85). The brainstem-spinal cord was dissected free and had the dura removed. It was then glued to an agar block and sliced (300 μm thick) with a vibrating microtome (Technical Products International Series 1000, St. Louis, MO, USA). Slices containing the hypoglossal motor pool were transferred to a 0.5 ml chamber and fixed to the chamber with a nylon grid for patch clamp electrophysiology experiments. The chamber was situated in a fixed stage microscope for imaging of the slice (FN1, Nikon Instruments Inc., Melville, NY, USA). Oxygenated aCSF was fed through the chamber with a peristaltic pump at ∼1.5 ml per minute.

Hypoglossal motoneurons in the rostral region were targeted as they are part of the buccal force pump for ventilation in anuran amphibians ^61,62^ and therefore play a role in breathing and other orofacial behaviors that are critical to survival ^42,43^. Hypoglossal motoneurons in the rostral part of the motor pool were identified anatomically based on their dorso-medial position relative to the 4^th^ ventricle and large cell body size (>20 μM diameter along the shortest axis) as previously described ^60^. Slices were imaged using a Hamamatsu ORCA Flash 4.0LT sCMOS camera (Hamamatsu Photonics, Hamamatsu City, Japan) interfaced with NIS elements software (Nikon). Once identified, motoneurons were approached by applying positive pressure through the glass microelectrode for patch clamp recording. Electrophysiological signals were amplified by an Axon Instruments 200B patch clamp amplifier and digitized with a Digidata 1550 digitizer (Molecular Devices). Microelectrodes were fabricated with a Sutter Instrument P-1000 micropipette puller (Novato, CA, USA) and had resistances of ∼5 MΩ when filled with the pipette filling solution (in mmol): 110 potassium gluconate, 2 MgCl_2_, 10 Hepes, 1 Na_2_-ATP, 0.1 Na_2_-GTP and 2.5 EGTA, pH ∼ 7.2 with KOH prepared with RNA-free water. All pipettes were filled with 7 µl of internal solution. As the motoneuron was approached, debris was cleared by the positive pressure and when an indentation was observed in the center of the cell body, positive pressure was removed, increasing the resistance between the microelectrode and the cell membrane. Negative pressure was then applied until a seal ≥1G was observed and was then broken using rapid negative pressure by mouth to obtain whole-cell access.

### Whole-cell current clamp recordings

Current clamp experiments were conducted using the same equipment previously described (Zubov et al., 2022). Firing rates were measured after we obtained whole-cell access, when motoneurons were injected with incremental steps of +50 pA (step duration = 500 ms) from -150 pA to 1300 pA.

### Whole-cell voltage clamp recordings

After the current clamp protocol, slices were exposed to aCSF with the voltage-gated Na^+^ channel blocker tetrodotoxin (TTX, 500 nM) during 5 min to record miniature excitatory postsynaptic currents (mEPSC) at -80 mV. Under these conditions, all mEPSCs are blocked by DNQX and are therefore generated by AMPA or kainate glutamate receptors ^25^. mEPSCs were analyzed for 60 s at the end of the 5 minute period. Assessments of putative “synaptic scaling” were performed exactly as described in Zubov, et al. ^31^, which follows the procedure original put forth by Turrigiano, et al. ^7^. This analysis was run on a sample of 60 successive mEPSCs per cell obtained in sampling period (22 cells for controls and hibernators).

After this period, we evoked AMPAR currents by focally applying 500 µM AMPA onto the soma of hypoglossal motoneurons. For this, we prepared a second borosilicate pipette with a tip diameter ∼5 μm filled with AMPA (500 μM) dissolved in aCSF. The second pipette was positioned ∼20 µm from the neuron of interest, which was driven by a Picospritzer II using 20 psi of pressure. The solution was focally applied until we could observe the maximum evoked current. To ensure each neuron responded maximally, we increased the duration of the puff from ∼10-200 ms. Once increasing the duration of the puff no longer increased from the previous duration, we set the puff duration to the minimum duration required to elicit the maximum current amplitude. After identifying the maximal AMPAR current, we exposed slices to aCSF with TTX (500 nM) and 1-naphthyl acetyl spermine (NASPM, 100 μM), a blocker of Ca^2+^-permeable AMPAR, for another 5-10 minutes, repeating the puffing with AMPA until no further decrease was observed. At least 3 stable currents agonist-induced were analyzed per treatment. After the end of this protocol, the hypoglossal cytoplasm was collected following the steps outlined in Pellizari, et al. ^63^ and stored in 200 μl of lysis buffer at -80°C for subsequent quantitative PCR (qPCR) analysis.

In another set of experiments, we aimed to measure the maximum and minimum evoked postsynaptic currents between 22 and 10 °C. The temperature from the aCSF bathing the slices was controlled using a bipolar in-line temperature controller (Model CL-100, Warner Instruments, Hamden, CT, USA) and monitored with a thermocouple positioned close to the slice. Before we accessed the cells as described above, we positioned a bipolar tungsten electrode (WE3ST0.1B10, Microprobes for Life Sciences, Gaithersburg, MD, USA) connected to an isolated pulse stimulator (A-M Systems, Model 2100, Carlsborg, WA, USA) at the solitary tract axons ^45^. Maximum current amplitude was determined by increasing the stimulus currents until asynchronous vesicle release was observed (EPSCs that trailed after the synchronous EPSC and were larger than spontaneous EPSCs). Under these conditions, the entire evoked EPSC is blocked by DNQX, indicating it is carried by AMPA or kainate-type glutamate receptors ^64^. The stimulus intensity was then reduced to maintain the largest possible synchronous EPSC amplitude without asynchronous activity and was elicited at 0.2 Hz. After this period, temperature was reduced by 2°C until 10 °C. Cells were then warmed to 22 °C. If current amplitude did not fully recover to the baseline level, cells were excluded from analysis. Spontaneous EPSC were analyzed for frequency and amplitude in the interval between stimulation events.

On a third set of experiments, we sought to measure the spontaneous excitatory postsynaptic currents (sEPSC) at minimum stimulation for evoked postsynaptic currents at 22, 16 and 10 °C. Using the same procedures described for the maximal stimulation experiments, EPSCs were first determined at 0.5 Hz and then stimulation intensity was decreased until no currents were observed. Then, stimulation was increased incrementally to allow 10 successive stimulation events until we found the minimum value that evoked one current with a consistent waveform that contained both failures and evoked responses. Since the minimum stimulation intensity was strongly affected by temperature (i.e., minimum stimulation at 22° would never evoke release at 16°C), EPSCs at minimum stimulation was determined using this protocol for each temperature separately. The EPSC were first measured for 30 s at 22 °C, after which the minimum stimulation was determined and EPSCs were recorded for 5 min. This protocol was repeated for 16 and 10 °C.

All patch clamp electrophysiology data were analyzed using either the Easy Electrophysiology data analysis program (Easy Electrophysiology Ltd., London, UK) or LabChart (v 8.0, ADInstruments, Oxford, UK). Firing rate was taken as the firing rate during 500 ms current injections and is presented as either maximal firing rate or the entire frequency vs. current relationship. Input resistance is taken as the change in voltage change from resting membrane potential during a 100 pA step. Maximal and minimal synaptic currents were assessed taking the difference between the peak and baseline current that proceeded stimulation. Probability of release was calculated as the number of successful stimulation events divided by the total number of stimulation events that occurred during the 5 min recording period.

## Drugs

Tetrodotoxin citrate (TTX), 1-naphthyl acetyl spermine (NASPM) and α-amino-3-hydroxy-5-methyl-4-isoxazolepropionic acid (AMPA) were acquired from Hello Bio (Princeton, NJ, USA).

## RNA isolation and real-time quantitative PCR

All methods for single-cell harvesting using the patch pipette, RNA extraction, cDNA synthesis, and qPCR were performed as described in Bueschke, et al. ^40^ and Pellizari, et al. ^63^. The only modifications was that harvesting process involved ∼7.5 µl of pipette filling solution containing the cytoplasm of the neuron following physiology recordings. These contents were then added to 200 µl of lysis buffer (Zymo Research) and then immediately frozen on dry ice. After all samples were collected, RNA was extracted from single-cell samples using Quick-RNA MicroPrep Kit (Zymo Research) according to the manufacture’s instructions. RNA was then reverse transcribed into the cDNA using SuperScript IV VILO in a 14 µL reaction according to the manufacturer’s instructions (ThermoFisher Scientific, Waltham, MA). After this step, 1 µL of cDNA was used as the template in a qPCR reaction to assess the expression of the housekeeping gene (18S ribosomal RNA (rRNA) and the other 13 µl was used as the template to preamplify target genes. For this, we used PerfeCTa PreAmp Supermix (Quanta BioDesign, Plan City, OH) to create a 20 µL reaction according to the manufacturer’s instructions. After preamplification, the contents of the tube were diluted 7.5-fold in nuclease-free water (150 µL final volume), which was then used as the template for real-time quantitative PCR of the target genes.

PCR primers for *Gria1*, *Gria2*, *Gria3*, *Gria4*, *Grik1,* and *Grik2* were designed based on sequences identified in the coding DNA sequence for *Aquarana catesbeiana*. Briefly, we used annotated amino acid sequences for our targets of interest from *Rana temporaria or Xenopus laevis* as a query in the *Aquarana catesbeiana* amino acid database. This produced hits with high amino acid sequence conservation. We then performed a reciprocal BLAST against the entire nonredundant protein database using hits from *Aquarana catesbeiana* to verify the identity of the target. Accession numbers were then used to identify the open reading frame in the CDS to design PCR primers.

Multiplex qPCR assays containing 3 targets each (Table 1) were each validated in-house with a series of four four-fold dilutions of brainstem cDNA using the thermal cycling conditions stated below. All assays used here produced efficiencies greater than 80%. All neurons were assessed for the expression of each target gene using qPCR. For the control gene, 18S rRNA, quantitative PCR was run in 10 µL reaction volumes containing 2.5 µM forward and reverse primers and followed the instructions of the 2X SYBR Green Mastermix (Applied Biosystems, ThermoFisher Scientific, Waltham, MA). Assays were run on 96-well plates on an Applied Biosystems QuantStudio 5 (Applied Biosystems, ThermoFisher Scientific, Waltham, MA) using the following cycling conditions according to the SYBR Green instructions: 50°C-2m, 95 °C-10 m, 95 °C-15 s, 60 °C-1 m. Following 40 cycles of PCR (95°C-15 s, 60°C-1 m), melt curves for all PCR products were acquired by increasing the temperature in increments of 0.3°C for 5 seconds from 60°C to 95 °C. *Gria1*, *Gria2*, *Gria3*, *Gria4*, *Grik1,* and *Grik2* were run using multiplexed probed-based assays. For this, we used the same primer concentration as described for SYBR Green assays, 312.5 nM reporter probes, and followed the instructions of the 5X PerfeCTa qPCR Toughmix mastermix (Quanta Bio). Absolute quantitation of transcript abundance was estimated through copy number standard curves as previous described ^9^.

**Table 1.**
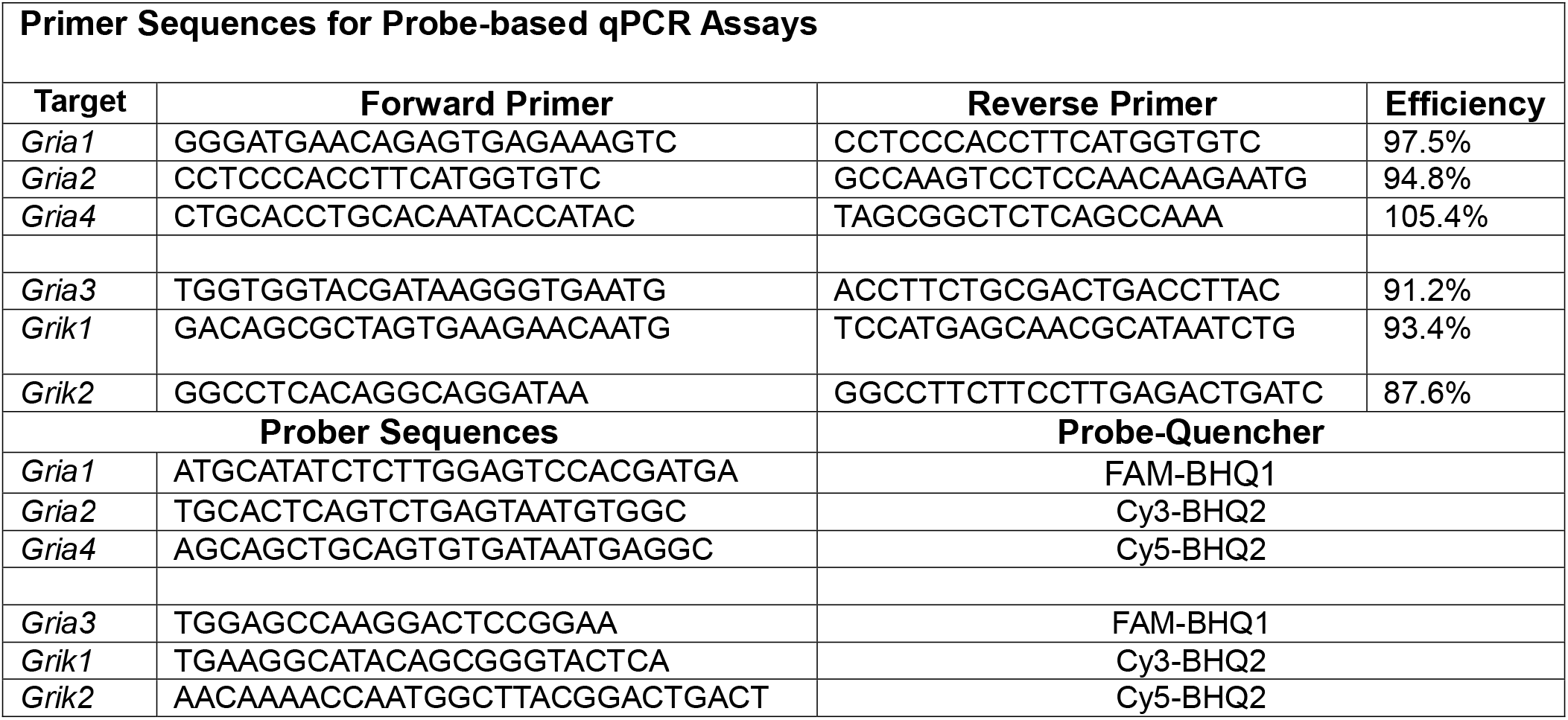
Primer and probe information for candidate genes.

## Statistics and data presentation

Data are presented as mean ± standard deviation or box and whisker plots, with single data points from individual experiments where it is most appropriate for visualization. When data sets were approximately normally distributed, parametric statistical tests were used. If data sets did not pass normality tests, non-parametric equivalents were used. When groups contained independent samples, a two-tailed unpaired t-test or one-way ANOVA was used. One-way and two-way ANOVAs were followed with Holm-Sidak posthoc tests. In cases where data sets with independent groups were not normally distributed, a Mann-Whitney test or a Kruskal-Wallis test was used when appropriate. Cumulative distributions were assessed with the Kolmogorov-Smirnov test, as is typically done for assessments of putative synaptic scaling ^7^. Null hypotheses were rejected when p < 0.05. All analyses were performed using GraphPad Prism (v9.4.1, San Diego, CA, USA).

## Acknowledgment

We would like to thank the National Institutes of Health (R01NS114514) and the National Science Foundation (2515635) for funding to JMS.

