## Supplemental Figure 1 for "Activity-dependent homeostatic synaptic plasticity widens the temperature range of synaptic transmission"

**Supplemental Materials**

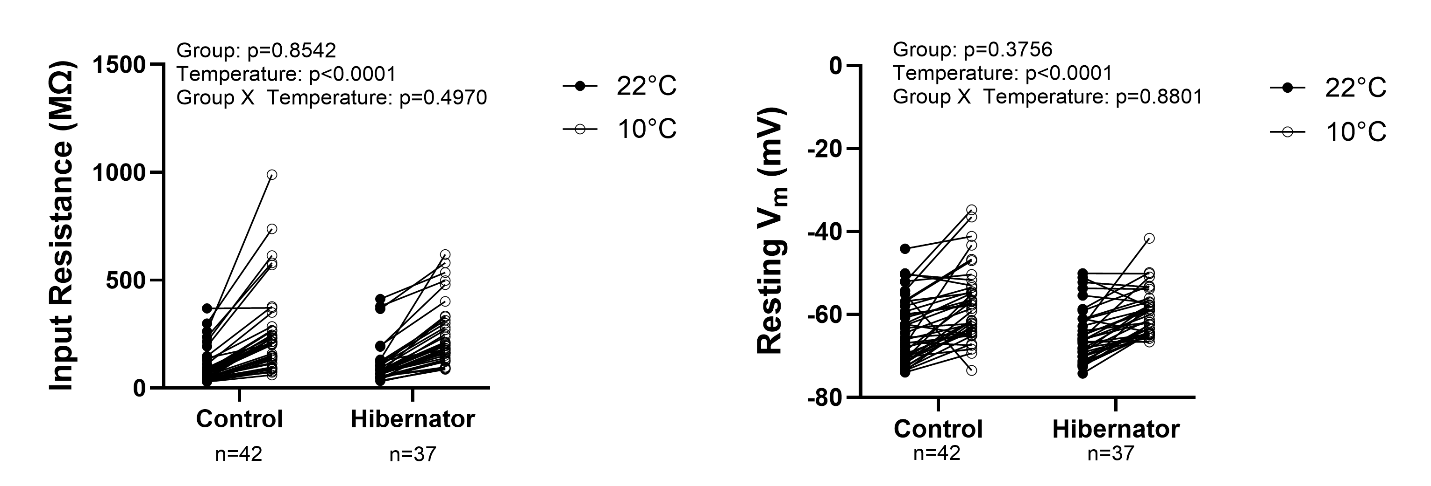

**Supplemental Figure 1 Temperature-dependent changes in passive membrane properties are the same in controls and hibernators.** Left shows input resistance in controls and hibernators before and after cooling to 10°C. Two-way ANOVA results indicate an expected increase in input resistance by cooling. However, there was no group effect or interaction between group and temperature. Right plot shows resting membrane potential in controls and hibernators before and after cooling to 10°C. Two-way ANOVA results indicate an expected depolarization of resting membrane potential by cooling. However, there was no group effect or interaction between group and temperature. Individual data points represent “before” and “after” cooling to 10°C. N=6 controls and N=6 hibernators were used to obtain the neuron “n” presented in the graphs.
